# Contrasting maternal and paternal genetic variation of hunter-gatherer groups in Thailand

**DOI:** 10.1101/201483

**Authors:** Wibhu Kutanan, Jatupol Kampuansai, Piya Changmai, Pavel Flegontov, Roland Schröeder, Enrico Macholdt, Alexander Hüebner, Daoroong Kangwanpong, Mark Stoneking

## Abstract

The Maniq and Mlabri are the only recorded nomadic hunter-gatherer groups in Thailand. Here, we sequenced complete mitochondrial (mt) DNA genomes and ~2.364 Mbp of non-recombining Y chromosome (NRY) to learn more about the origins of these two enigmatic populations. Both groups exhibited low genetic diversity compared to other Thai populations, and contrasting patterns of mtDNA and NRY diversity: there was greater mtDNA diversity in the Maniq than in the Mlabri, while the converse was true for the NRY. We found basal uniparental lineages in the Maniq, namely mtDNA haplogroups M21a, R21 and M17a, and NRY haplogroup K. Overall, the Maniq are genetically similar to other negrito groups in Southeast Asia. By contrast, the Mlabri haplogroups (B5a1b1 for mtDNA and O1b1a1a1b and O1b1a1a1b1a1 for the NRY) are common lineages in Southeast Asian non-negrito groups, and overall the Mlabri are genetically similar to their linguistic relatives (Htin and Khmu) and other groups from northeastern Thailand. In agreement with previous studies of the Mlabri, our results indicate that the Malbri do not directly descend from the indigenous negritos. Instead, they likely have a recent origin (within the past 1,000 years) by an extreme founder event (involving just one maternal and two paternal lineages) from an agricultural group, most likely the Htin or a closely-related group.

## Introduction

The indigenous peoples of Southeast Asia (SEA) are often referred to as “Negritos” because of their differences in phenotypic appearances, i.e. on average shorter stature, frizzier hair, and darker skin color^1–2^. The phenotypic resemblances of SEA negritos and African pygmies initially suggested a separate origin in Africa, although convergent evolution now seems more likely (for reviews see Détroit et al.^3^ and Stock^4^). Hunting-gathering was the mode of subsistence traditionally practiced by most SEA negritos, whose communities are distributed in three regions: Andaman Islands, Peninsular Thailand and Malaysia, and the Philippines^5^. Several genetic studies have been conducted in some of these relic populations and indicated deep ancestry^6–13^ which could link these people with the ancestral groups who migrated from Africa into SEA during the late Pleistocene^14^. However, the hunter-gatherer groups of Thailand are relatively under-studied compared to those from other SEA regions.

There are two hunter-gatherer groups in Thailand, the Maniq and the Mlabri. The Maniq, also known as Sakai or Kensui, have a census size of ~250 individuals^15–16^ and are considered to be related to the Semang of Malaysia^17^. For example, the Maniq language belongs to the northern Aslian group of Austroasiatic languages, as do the Semang languages^18^, although this most likely reflects a later introduction by agriculturalists^19–20^. A previous study of sequences of the first hypervariable region (HVR) of the mtDNA control region reported a relatively large divergence of the Sakai (Maniq) from southern Thailand from other Thai groups^21^.

The other hunter-gatherer group, the Mlabri, are characterized by a nomadic traditional lifestyle, in which they construct shelters from the leaves of banana trees that they move out when the banana leaves turn yellow. Their neighbors call them “Phi Tong Luang” which means the “Spirits of the Yellow Leaves”. Although they do not exhibit the typical “negrito” phenotypes, their mode of subsistence is hunting and gathering^22–23^. This, however, is disputed by Waters (2005)^24^ who believed that the Mlabri have had contact and trade with neighboring ethnic groups, and thus they are not hunter-gatherer in the strict sense. The conflicting theories as to how they sustain their livelihood make it interesting to study their origin and lineages. With a census size of only ~130 individuals^16^, the Mlabri live in a mountainous area of northern Thailand and speak a language belonging to the Khmuic branch of Austroasiatic languages. Khmuic languages are also spoken by their neighbors, Htin (of Mal and Pray subgroups) and Khmu, who are also the other hilltribes in northern Thailand. Preceding genetic studies reported extremely reduced genetic diversity in the Mlabri and suggested a cultural reversion from agriculture to hunting and gathering^25^ and genetic affinities between the Mlabri and the Mal, a Htin subgroup^26^.

Comparisons of variation in maternally-inherited mtDNA and paternally-inherited, non-recombining portions of the Y chromosome (NRY) have been useful in illuminating differences in the female vs. male demographic history of human populations^27–28^. However, such comparisons are complicated by the different molecular methods that have been typically employed to assay variation (e.g., HVR sequencing for mtDNA vs. genotyping of SNPs and/or STR loci for the NRY). Advances in next-generation sequencing make it feasible to obtain mtDNA and NRY sequences that allow directly comparable, unbiased inferences about female vs. male demographic history^29–30^. Here, we present and analyze complete mtDNA genome sequences and ~2.364 Mbp of NRY sequence from the Maniq and the Mlabri (Fig. 1). These two groups have very distinct mtDNA and NRY lineages and contrasting patterns of variation. Overall, our results support an affiliation of the Maniq with other SEA negrito groups, and a recent origin of the Mlabri lineages.

**Figure 1.**
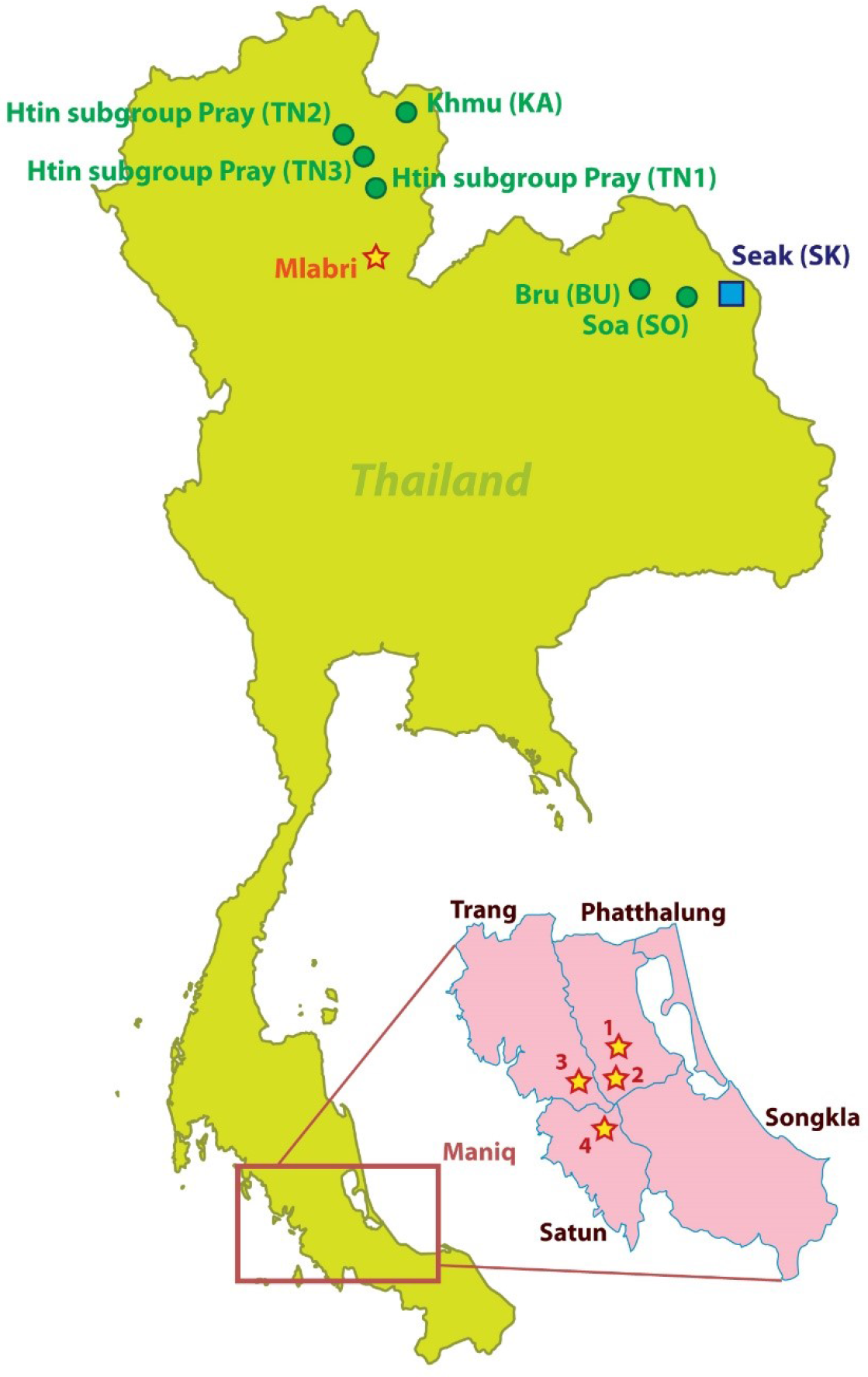
Map of the Mlabri and Maniq sampling locations (stars) and the Thai populations selected for NRY comparisons (comparative mtDNA data are from previous studies^31,32^). Further details concerning the populations are provided in Table S1 and S2. There are four Maniq collection sites; Pabon District (site 1) and Kong-Ra District (site 2), Phatthalung Province; Palien District, Trang Province (site 3); and La-Ngu District, Satul Province (site 4).

## Results

### Mitochondrial DNA

We generated 29 complete mtDNA sequences (18 from Mlabri and 11 from Maniq) with mean coverages ranging from 277X to 1342X. Extraordinarily low mtDNA diversity was observed in the Mlabri (*h* = 0.29 ± 0.12; MPD = 0.29 ± 0.32; *π* = 0.00002 ± 0.00002; *S* = 1) with just one polymorphic site (nt 43) defining two haplotypes. With 5 haplotypes defined, the Maniq showed higher diversity values (*h* = 0.82 ± 0.08; MPD = 29.85 ± 14.16; *π* = 0.0018 ± 0.00097; *S* = 64) than the Mlabri. However, both of them have lower genetic diversity values than other populations from Thailand (Figure S1)^31–32^.

The two different mtDNA sequences found in the Mlabri belong to haplogroup B5a1b1 and have a coalescent age of ~735.48 ya (95% HPD: 1.71 – 2321.88; Figure S2). The overall coalescent age of B5a1b1 is ~13.99 kya, which was similar to our previous study^32^. The network analysis (Fig. 2) also supports a single maternal ancestor for the Mlabri. The average number of mutations among the 18 Mlabri sequences is 0.29; with a mutation rate of one mutation in every 3,624 years^33^, this suggests an age for the Mlabri mtDNA variation of ~1,000 years, close to the coalescence age estimate. Overall, the mtDNA results are in keeping with previous analyses suggesting a single maternal ancestor for the Mlabri and a recent founding age within the past 1000 years^25^.

**Figure 2.**
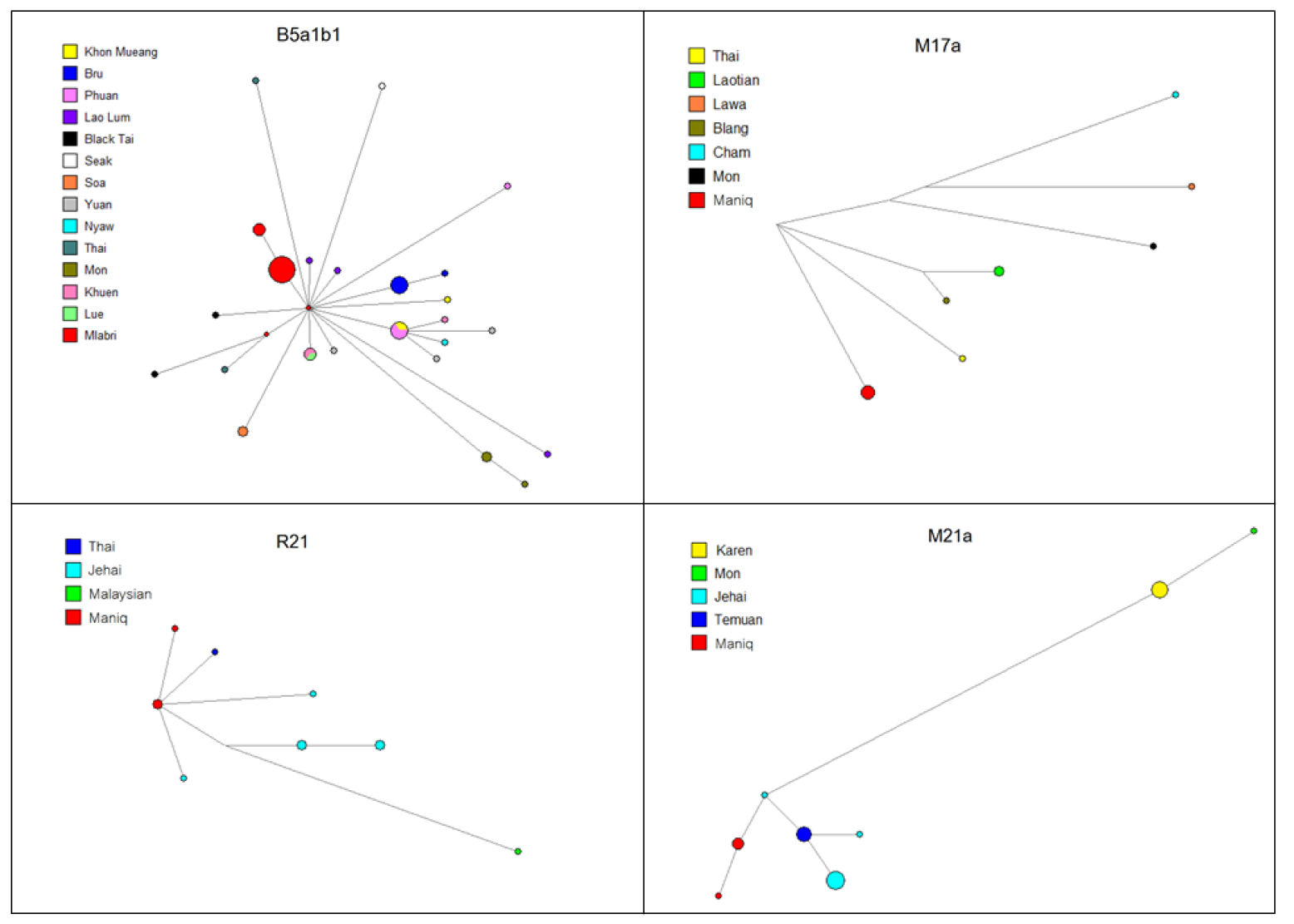
Networks of mtDNA haplogroups: B5a1b1 is found in Mlabri and R21, M21a and M17a are found in Maniq.

By contrast, there are three basal haplogroups (M17a, M21a and R21) observed in the Maniq. When combined with sequences from other populations (Table S1), the respective estimated ages of M17a, M21a and R21 are ~23.52, ~18.44 and ~8.83 kya (Figure S3). The ages of M17a and M21a are much older than our previous estimations in a Thai dataset^11,–32^ while R21 is newly-reported in populations from Thailand. The networks and MCC trees of M21a and R21 indicate a close relatedness of the Maniq sequences with those from the Jehai (an aboriginal Malaysian negrito group), while the Maniq M17a sequences cluster with sequences from the Laotian and Austroasiatic-speaking Blang (Fig. 2; Figure S3).

The MDS plot based on phiST distances indicates that the studied groups have large genetic distances from each other and from most other populations (Fig. 3a). In the NJ tree (Fig. 3b), the Mlabri do not show any close relationship with other SEA hunter-gatherer or negrito groups, but instead are closest to other northeastern Thai populations (Seak, Soa and Bru). In contrast, the Maniq cluster with populations from West Malaysia, and are closest to the Jehai, one of the Semang negrito groups of Malaysia (Fig. 3b).

**Figure 3.**
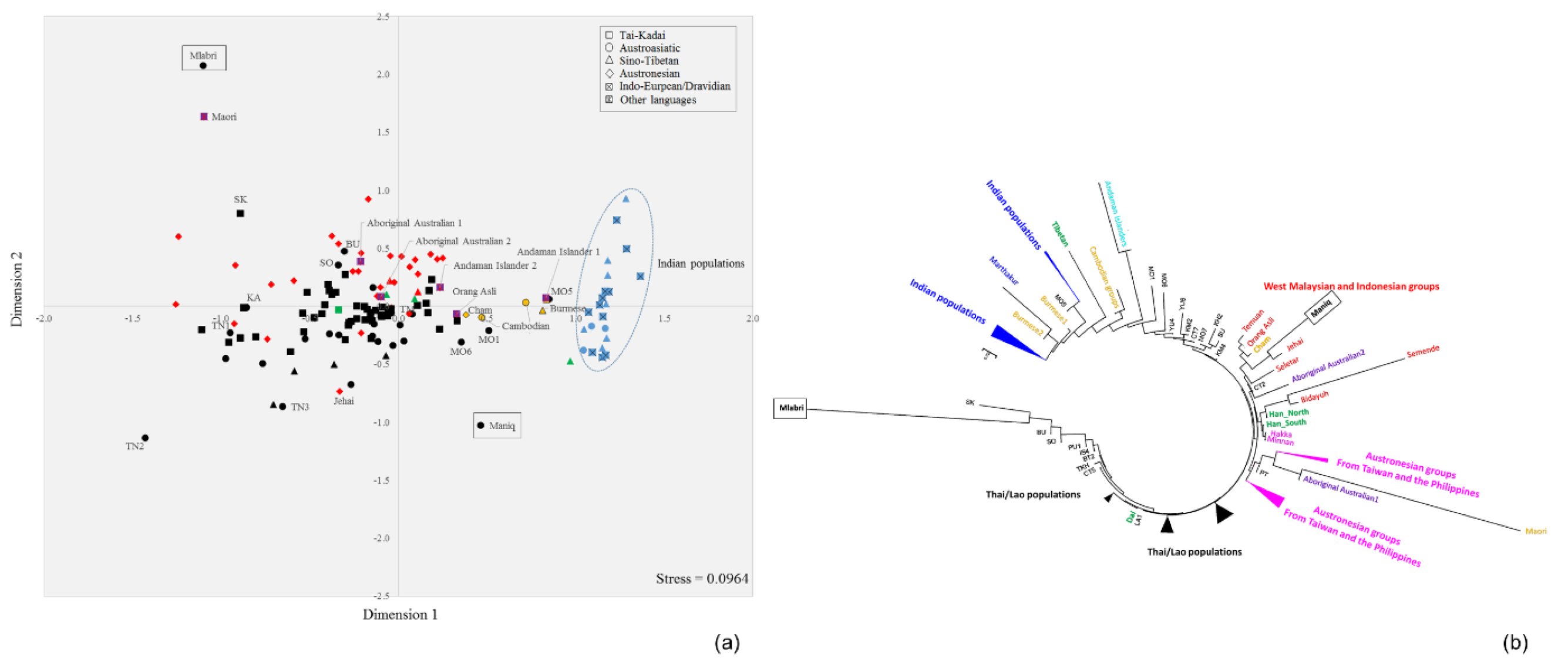
The MDS plot (a) and NJ tree (b) based on the mtDNA *Φ_st_* distance matrix for 142 populations from Asia, New Zealand and Australia. Population details are provided in Table S1. TN1 is Htin subgroup Mal and TN2 and TN3 are Htin subgroup Pray. KA, SO, SK and BU are Khmu, So, Seak and Bru, respectively. Black indicates Thai/Lao populations; red indicates populations from Indonesia and Malaysia; green indicates populations from China.

### Y chromosome

We generated 14 sequences (10 from the Mlabri and 4 from the Maniq) for ~2.364 Mb of the NRY, with mean coverages ranging from 10X to 25X. In contrast to the mtDNA results, the Mlabri have higher NRY diversity (six different sequences; *h* = 0.7778 ± 0.1374; MPD = 17.46 ± 8.94; *π* = 7.39 × 10^−6^; *S* = 37) than the Maniq, which show just one haplotype (i.e., all four sequences were identical). It should be noted that these four unrelated sequences were collected from three different locations (sites 1, 3 and 4) (Fig. 1).

Among six Y chromosomal haplotypes in ten Mlabri males, an MCC tree shows two clades: one comprises five haplotypes of haplogroup O1b1a1a1b which diverged ~1.43 kya and the other is a single haplotype, found in five individuals, classified to haplogroup O1b1a1a1b1a1 (Figure S4), suggesting two paternal lineages for the Mlabri. When we combine the Y sequences of haplogroup O1b1a1a1b and its sublineages retrieved from previous studies (Table S2), the newly reported age of haplogroup O1b1a1a1b is ~6.89 kya with a signal of population expansion. The MCC tree shows that the O1b1a1a1b sequences of the Mlabri are clustered with sequences from the Htin subgroup Mal, while the O1b1a1a1b1a1 sequences are clustered with sequences from the Khmu and the Htin subgroup Pray (Figure S4). The network of haplogroup O1b1a1a1b shows an overall star-like structure, suggesting lineage expansion, and further supports two paternal lineages for the Mlabri (Fig. 4). The NRY sequence of the four unrelated Maniq males belongs to haplogroup K, and when compared to the two haplogroup K2b1 sequences^30^, the estimated coalescent age is ~41.10 kya (Figure S5). The Maniq sequence thus adds to a small set of basal Y chromosomes sequences.

**Figure 4.**
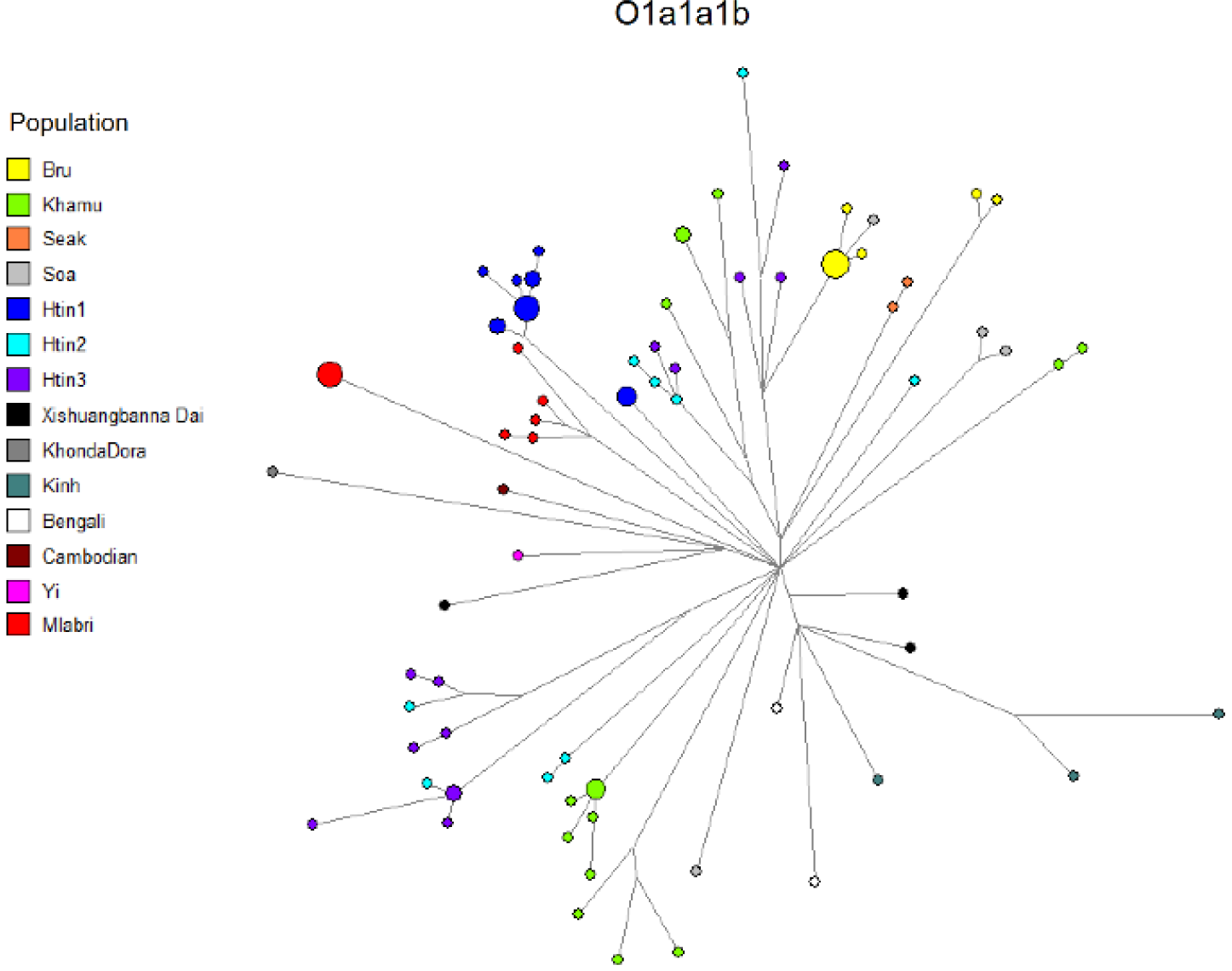
Network of NRY haplogroup O1a1a1b sequences.

The Mlabri NRY sequences cluster with other northern Thai populations in the MDS plot and NJ tree (Fig. 5a and 5b), particularly with their linguistic relatives, Khmu and the Htin subgroup Pray. Although the Maniq are highly diverged from other populations due to their very low diversity, they show closest affinity with Papuan, island SEA (ISEA), and South Asian populations (Fig. 5b).

**Figure 5.**
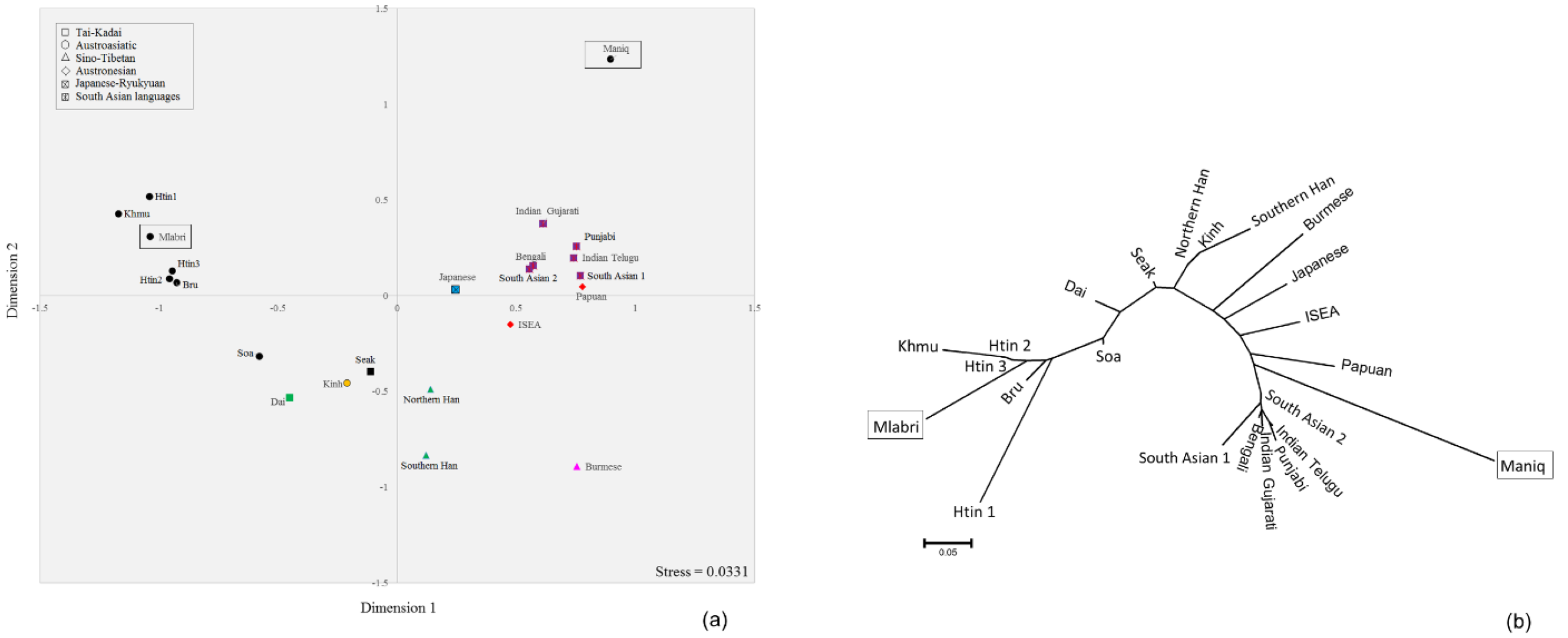
The MDS plot (a) and NJ tree (b) based on the Y chromosomal *Φ_st_* distance matrix for 23 populations from Asia and Oceania. Black circles indicates Thai populations. Population details are provided in Table S2.

## Discussion

We here report and analyze sequences of mtDNA genomes and ~2.364 Mbp of the NRY from the Mlabri and Maniq, two enigmatic hunter-gatherer groups from Thailand. The large genetic divergence of the Mlabri and the Maniq from each other and from the other populations probably reflects genetic drift associated with isolation and very small population sizes. Notably, we observed contrasting patterns of variation in mtDNA vs. the NRY: mtDNA diversity is higher in the Maniq and lower in the Mlabri, while the reverse is true for the NRY.

The SEA negritos are of interest because they are characterized by basal genetic lineages that likely reflect the earliest dispersal of modern humans from Africa^6,34^. However, there is also evidence that at least some negrito groups have interacted and mixed with non-negrito groups^10–12,35^. Among the three basal mtDNA haplogroups (M21a, M17a and R21) observed in the Maniq (Fig. 2), two (M21a and R21) have been previously reported in negrito groups from Malaysia^9,13^. MtDNA lineages of the Maniq (also known as the Kensui) were mentioned as unpublished data in a previous study^13^, which also found M21a and R21 to be common lineages, along with several minor SEA-specific haplogroups (B5a, F1a1a, M7c3c and N9a6a), indicating some SEA influences. In contrast, we did not detect any SEA-specific mtDNA haplogroups (B, F and M7) in the Maniq, but instead we found M17a, which has been reported at low frequency in MSEA^32,36^ and ISEA^37^. Moreover, we detected Y chromosomal haplogroup K in the Maniq, which occurs elsewhere in populations of Papua New Guinea^38–39^, the Philippines^28^ and Australia^43^. Haplogroup K is an ancient lineages that is dated here to ~41.10 kya, which is younger than previous estimates in Aboriginal Australians (~48.40 kya)^40^. Overall, the Maniq display basal genetic lineages and closest genetic affinities with indigenous groups in Malaysia or ISEA, supporting a common ancestry of the Maniq with other SEA negrito communities. The reduced genetic diversity (and population size) of the Maniq probably occurred as a consequence of agricultural expansions to this region. The archaeological evidence in southern Thailand suggests that the Neolithic period of the Malay Peninsula started around 4 kya^19^ and after the expansion of agriculturalists who also brought Austroasiatic languages to this region, the Proto-Aslian groups were fragmented^20^, probably leading to gradual isolation.

The Mlabri present a different picture. Their mtDNA and NRY haplotypes are closely related to those of neighboring agricultural groups, rather than belonging to basal haplogroups that are characteristic of typical hunter-gatherer groups. Overall, the mtDNA and NRY results suggest a severe founder event in the Mlabri ancestors, involving just one mtDNA lineage and two NRY lineages that occurred ~1000 ya. These results are in good agreement with previous studies that suggested that the Mlabri originated from an agricultural group and reverted to hunting-gathering^25^. We further attempted to identify the closest living relatives of the Mlabri, as these might be their source population. We found close genetic relationships in the NRY lineages between the Mlabri and various Htin subgroups as well as the Khmu, in agreement with an oral tradition, which indicates that the Htin are the ancestors of the Mlabri^25^. A shared common ancestor of these Khmuic-speaking groups might be another explanation. However, the Mlabri mtDNA lineage is most closely related to the Seak, Soa and Bru groups from northeastern Thailand (Fig. 3). Historical evidence indicates that the Mlabri, Soa, Seak and Bru in Thailand migrated from the present-day area of Laos about 100-200 ya^23^. Although each group migrated independently, their close relatedness might reflect maternal gene flow among various groups in Laos. However, the Mlabri stand out among these groups in apparently having just one maternal ancestor (Fig. 2).

A limitation of our study is that the sample sizes are quite small, which reflects the difficulty in contacting these elusive and enigmatic groups. Nonetheless, a rigorous participant recruitment procedure, and overall agreement with previous studies, suggests that our results are indeed representative of these small populations. Notably, the Mlabri are probably not indigenous SEA hunter-gatherers, but rather originated recently from an agricultural group via an extreme founder event, reflected in their one maternal and two paternal ancestral lineages. The Htin and Khmu are their potential ancestors. By contrast, the Maniq are likely to be indigenous SEA hunter-gatherers whose ancient ancestry is not mixed with non-negrito populations; further study of additional Maniq samples from the other bands as well as autosomal data would provide more information on the history and relationships of this group.

## Methods

### Samples

Genomic DNA samples from 18 Mlabri who live in a village located in Wieng Sa District, Nan Province, Northern Thailand were obtained from a previous study^41^. Because the census size of the Mlabri is about 130 individuals, we collected all possible unrelated samples which could be assumed to be representative of this group. The Maniq still lead a highly mobile way of life in the forested Banthad Mountain of three southern Thai Provinces, i.e. Patthalung, Trang and Satun. They live in about 9 small bands or groups, with 8-50 individuals per group, and a total census size of about 250 individuals. All groups are quite closely related, and there is constant movement of family members between groups^15,17^. We obtained saliva samples from 11 Maniq from four different bands in three provinces, i.e. Pabon District (site 1: 3 samples) and Kong-Ra District (site 2: 1 sample), Phatthalung Province; Palien District, Trang Province (site 3: 2 samples); and La-Ngu District, Satun Province (site 4: 5 samples) (Fig. 1). Informed consent was obtained prior to the collection of Maniq saliva samples, and all volunteers were interviewed by interpreters and coordinators to screen for subjects unrelated for at least two generations. We extracted DNA using the QIAamp DNA Midi Kit (Qiagen, Germany) according to manufacturer’s directions. Research on human subjects was approved by Khon Kaen University, and the Ethics Commission of the University of Leipzig Medical Faculty.

### Sequencing

#### mtDNA

Twenty-nine complete mtDNA sequences (18 Mlabri and 11 Maniq) were generated from genomic libraries prepared with double indices and enriched for mtDNA as described previously^42–43^; the libraries were sequenced on the Illumina Hiseq 2500. MtDNA consensus sequences were obtained as described by Arias-Alvis et al.^44^ with minor modifications. We used Bustard for Illumina standard base calling with a read length of 76 bp. We manually checked sequences with Bioedit (www.mbio.ncsu.edu/BioEdit/bioedit.html) and aligned the sequences to the Reconstructed Sapiens Reference Sequence (RSRS)^45^ using MAFFT 7.271^46^.

#### Y chromosome

We designed a capture probe set for the NRY following a previous study^47^ and using genome sequences incluldedin SGDP Panel A^48^ and Panel B^49^. Sites were kept that were included in the UCSC table browser *CRG Align* 100 mappability track of the human reference genome *hg19* (9,655,499 bp). In order to obtain confident Y haplogroup calls later, we selected regions with at least one ISOGG SNP (version 8.78) and then applied the following filter criterion: a region was removed when its length was lower than 500 bp and its repeat count of 24-mers was higher than 10 in an average window of 120 bp using the *CRG Align 24* mappability track from the UCSC table browser. A total of 2,264 regions with a combined length of 2,364,049 bp were included into the capture probe set, which included 2,071 ISOGG SNPs (version 11.144).

Genomic libraries were prepared for each sample using a double indices scheme^48^. We enriched the libraries for the NRY regions via in-solution hybridization-capture^49^ using our probe set and the Agilent Sure Select system (Agilent, CA, USA) and sequenced on the Illumina HiSeq 2500 platform, generating paired-end reads of 125 bp length. Standard Illumina base-calling was performed using Bustard. We trimmed Illumina adapters and merged completely overlapping paired sequences using leeHOM^50^ and de-multiplexed the pooled sequencing data using deML^51^.

The sequencing data were aligned against the human reference genome *hg19* using BWA’s *aln* algorithm^52^ and all sequence pairs that aligned to the NRY region which are previously defined were retained^47^. Duplicate reads were removed using PicardTools *MarkDuplicates*, indel realignment was performed using GATK *IndelRealigner*^53^ and base quality were re-calibrated using GATK *BaseRecalibrator*. We identified single nucleotide variants (SNVs) using GATK *UnifiedGenotyper* v3.3-0 across all samples simultaneously setting the parameter *ploidy* to 1 and using DBSNP build 138 as a prior position list. The identified SNVs were further filtered as previously described^54^. We imputed all samples with missing genotype information at any of these variant sites using BEAGLE^55^.

### Statistical analyses

Haplogroup assignment was performed by Haplogrep^56^ for mtDNA and yHaplo^57^ for the NRY. Summary statistics for the studied populations and genetic distances (*Φ_st_*) among the studied and compared populations were computed by Arlequin 3.5.1.3^58^. Comparative data for the mtDNA genome sequences were selected from the literature encompassing East Asia^31–32,36,59–64^, Island Southeast Asia^12–13,34,65–67^, Taiwan^68^, India^69^, Andaman Islands^6,70^, Australia^71–73^ and New Zealand^74^. Further information on the populations used for mtDNA comparison are listed in Table S1. Overlapping NRY sequences of relevant populations from Thailand (Htin, Khmu, Soa, Bru and Seak) (unpublished data), East Asia, Oceania, South Asia and also Indian groups from the United Kingdom and the United States of America^30,75–76^ were included for comparison; additional details are provided in Table S2.

The *Φ_st_* distance matrices of both mtDNA and the NRY were used to construct multi-dimentional scaling (MDS) plots and Neighbor Joining (NJ) trees^77^ by STATISTICA 13.0 (StatSoft, Inc., USA) and MEGA 7.0^78^, respectively. Median-joining networks^79^ of haplogroups without pre- and post-processing steps were performed by Network (www.fluxus-engineering.com) and visualized in Network publisher 1.3.0.0.

BEAST 1.8.0 was used to construct maximum clade credibility (MCC) trees by haplogroup, based on Bayesian Markov Chain Monte Carlo (MCMC) analyses^80^. We ran jModel test 2.1.7^81^ for selecting the suitable models in each run during the creation of the input file of BEAST via BEAUTi v1.8.0^80^. For mtDNA, the BEAST calculation was done with the data partitioned between coding and noncoding regions with mutation rates of 1.708 × 10^−8^ and 9.883 × 10^−8^, respectively^33^. For the NRY, we used a mutation rate of 8.71 × 10^−10^,^82^ and the BEAST input files were modified by an in-house script to add in the invariant sites found in our dataset. After BEAST runs, we used Tracer 1.6 to check the results and TreeAnnotator v1.6.2 and FigTree v 1.4.0. to assemble and draw the Bayesian MCC trees, respectively. The Bayesian MCMC estimates (BE) and credible intervals (CI) of haplogroup coalescent times were calculated using the RSRS for rooting the mtDNA tree and African sequences belonging to haplogroup B2 (GS000035246-ASM) for rooting the NRY tree^30^.

## Acknowledgements

We would like to thank the participants and the people involved with participant recruitment. We thank Prof. Murray Cox for discussion and valuable suggestions. W.K. and M.S. were supported by the Max Planck Society. WK was also funded by the Thailand Research Fund (Grant No. MRG5980146). P.F. was supported by the Institution Development Program of the University of Ostrava and by the Rector’s award of the University of Ostrava. P.C. was supported by the grant no. 0924/2016/ŠaS from the Statutory City of Ostrava and by the grant no. 01211/2016/RRC “Strengthening international cooperation in science, research and education” from the Moravian-Silesian Region.

## Author Contributions

W.K. and M.S. conceived and designed the experiment, provided funding, and drafted the manuscript; W.K. and R.S. conducted sequencing; J.K., P.C., P.F. and D.K. collected the samples and extracted DNA; W.K., A.H. and E.M. analysed the results; All authors reviewed the manuscript.

## Additional information

### Data availability

GenBank accession numbers will be provided upon acceptance for publication.

